# Auto-Immunoproteomics Analysis of COVID-19 ICU Patients Revealed Increased Levels of Autoantibodies Related to Male Reproductive System

**DOI:** 10.1101/2022.02.09.479669

**Authors:** Frank Schmidt, Houari B. Abdesselem, Karsten Suhre, Muhammad U. Sohail, Maryam Al-Nesf, Ilham Bensmail, Fathima Mashod, Hina Sarwath, Joerg Bernhardt, Ti-Myen Tan, Priscilla E Morris, Edward J. Schenck, David Price, Nishant N. Vaikath, Vidya Mohamed-Ali, Mohammed Al-Maadheed, Abdelilah Arredouani, Julie Decock, Jonathan M. Blackburn, Augustine M.K. Choi, Omar M. El-Agnaf

**Author notes:** These authors contributed equally to the work (Frank Schmidt, Houari B. Abdesselem). Corresponding Authors: Frank Schmidt, Address: Proteomics Core, Weill Cornell Medicine - Qatar, Doha, Qatar, E-Mail, Omar El-Agnaf, Address: Neurological Disorders Research Center, QBRI, HBKU, Qatar Foundation, Doha, Qatar.

## Abstract

The role of autoantibodies in coronavirus disease (COVID-19) complications is not yet fully understood. The current investigation screened two independent cohorts of 97 COVID-19 patients (Discovery (Disc) cohort from Qatar (n = 49) and Replication (Rep) cohort from New York (n = 48)) utilizing high-throughput KoRectly Expressed (KREX) immunome protein-array technology. Autoantibody responses to 57 proteins were significantly altered in the COVID-19 Disc cohort compared to healthy controls (P ≤ 0.05). The Rep cohort had altered autoantibody responses against 26 proteins compared to non-COVID-19 ICU patients that served as controls. Both cohorts showed substantial similarities (r^2^ = 0.73) and exhibited higher autoantibodies responses to numerous transcription factors, immunomodulatory proteins, and human disease markers. Analysis of the combined cohorts revealed elevated autoantibody responses against SPANXN4, STK25, ATF4, PRKD2, and CHMP3 proteins in COVID-19 patients. KREX analysis of the specific IgG autoantibody responses indicates that the targeted host proteins are supposedly increased in COVID-19 patients. The autoantigen-autoantibody response was cross-validated for SPANXN4 and STK25 proteins using Uniprot BLASTP and sequence alignment tools. SPANXN4 is essential for spermiogenesis and male fertility, which may predict a potential role for this protein in COVID-19 associated male reproductive tract complications and warrants further research.

**Significance Statement:** Coronavirus disease (COVID-19), caused by the SARS-CoV-2 virus, has emerged as a global pandemic with a high morbidity rate and multiorgan complications. It is observed that the host immune system contributes to the varied responses to COVID-19 pathogenesis. Autoantibodies, immune system proteins that mistakenly target the body’s own tissue, may underlie some of this variation. We screened total IgG autoantibody responses against 1,318 human proteins in two COVID-19 patient cohorts. We observed several novel markers in COVID-19 patients that are associated with male fertility, such as sperm protein SPANXN4, STK25, and the apoptotic factor ATF4. Particularly, elevated levels of autoantibodies against the testicular tissue-specific protein SPANXN4 offer significant evidence of anticipating the protein role in COVID-19 associated male reproductive complications.

## Introduction

Coronavirus disease (COVID-19), caused by novel SARS-CoV-2 virus, has emerged as global pandemic with severe complications and high morbidity rate. The disease manifests a wide range of clinical symptoms, which are exacerbated by overactive and malfunctioning immune system of the host. Despite extensive research on innate and adaptive immune responses in COVID-19, little is known about the role of autoantibodies on disease progression and severe complications.

Infection with the SARS-CoV-2 causes a variety of symptoms, with most cases being moderate or asymptomatic, and only a smaller proportion advancing to more severe state of COVID-19 disease^1^. Many questions about the COVID-19 pathophysiology remain open, particularly why some people develop severe disease symptoms while others remain asymptomatic.

Acute respiratory distress syndrome (ARDS) affects a small percentage of patients, whereas others experience persistent lung damage and multi-organ illness that lasts months, even after the virus has been eliminated from the body^2^. High expression of angiotensin-converting enzyme 2 (ACE2) receptors in several organs of the body extends infection beyond respiratory tract, resulting in complex multiorgan complications^3^. ACE2 receptors are highly expressed in the male reproductive system, demonstrating the involvement of SARS-CoV-2 in male fertility, which is one of the unexplained manifestations of COVID-19^4^.

Autoantibodies have been identified in significant proportion of COVID-19 hospitalized patients with positive correlation with immune responses to SARS-CoV-2 proteins^5^. Several studies observed significant rise in a diverse range of autoantibodies against immunomodulatory proteins, a- and w-interferons, cardiolipin and prothrombin during antiviral responses in severely ill COVID-19 patients^6, 7, 8, 9, 10, 11^. Particularly, autoantibodies against immune-related signaling proteins were found to contribute to COVID-19 pathogenesis by antagonizing the function of the innate immune system^12^. Although there have been some reports on disease-modifying autoantibody responses, the immunological and clinical consequences of autoantibodies in COVID-19 are yet to be fully understood. Here, we therefore screened total IgG autoantibody responses against 1,318 human proteins in COVID-19 patients using KREX immunome protein-array technology. Sengenics KREX technology employs full-length, naturally folded proteins that allow maximum epitopes binding to discover autoantibody biomarker proteins^13^. The quantitative signal measured on the arrays for each autoantibody-autoantigen pair is directly proportional to the autoantibody concentration in the blood with higher autoantibody titres to these proteins simplistically implying higher autoantigen concentrations in COVID-19 patients compared to controls, albeit the correlation is non-linear.

Autoantibody-based precision immuno-profiling has previously been shown to aid discovery of biomarkers of immune-related adverse events, as well as therapeutic prediction of drug response^14^. In the present study, by utilizing a broad array-based immunoproteomics strategy that simultaneously quantifies autoantibody responses across multiple organ systems in ICU COVID-19 patients and post recovery cohort, we aimed to better identify novel markers of comorbidities in COVID-19 patients. We identified a number of novel markers in COVID-19 patients that are also associated with male fertility, such as the sperm protein SPANXN4^15^, the androgenic kinase STK25^16, 17^, the apoptotic factor ATF4^18^, the calcium channel regulator protein kinase PRKD2^19^, and the multivesicular protein CHMP3^20^.

## Methods

### Study design, samples collection and processing and ethics

We used blood samples and clinical data of patients from two independent COVID-19 cohorts to conduct a comprehenisve anlaysis of autoantibodies using novel KREX technology.

#### Discovery (Disc) cohort

The Disc cohort included forty-nine COVID-19 patients from Qatar who were admitted to Hamad Medical Corporation hospitals. All recruited patients had confirmed SARS-CoV-2 positive RT-PCR results of sputum and throat swabs. All patients had severe COVID-19 disease (WHO guideline)^21^ and were admitted to intensive care unit (ICU). Peripheral blood was collected within five to seven days of admission and processed into plasma and serum, which were stored at –80°C, until further analysis. Ethical approval for this cohort was obtained from the Hamad Medical Corporation Institutional Review Board Research Ethics Committee (reference MRC-05-003), and Qatar Biomedical Research Institute-Institutional Review Board (Reference QBRI-IRB 2020-06-19).

#### Healthy controls

Age and gender matched healthy volunteers (n=48) with no prior COVID-19 infection history and with normal oxygen saturation and vital signs were used as controls. The Anti-Doping Laboratory-Qatar recruited them for blood collection. Individuals with medical history or with cognitive disabilities were excluded. All participants (patients and controls) provided written informed consent prior to enrolment in the study.

#### Replication (Rep) cohort

The replication cohort consisted of forty-eight adult patients who were admitted to the ICU of New York-Presbyterian Hospital (NYP)/Weill Cornell Medical Center (WCMC) from March to April 2020. All patients were RT-PCR confirmed SARS-CoV-2 positive and displayed ARDS or pneumonia symptoms. The cohort is part of the Weill Cornell Biobank of Critical Illness, a registry which attempts to recruit and enroll all patients being admitted to WCMC ICU for clinical investigations. The WCMC COVID Institutional Data Repository (COVID-IDR), a manually abstracted registry of COVID-19 patients that was developed to record patient demographics and allied health parameters. Laboratory parameters, ventilation records, respiratory variables and vital signs were recorded and documented at Weill Cornell-Critical Care Database for Advanced Research (WC-CEDAR)^22^. The processes for recruiting patients, collecting data, and processing samples had all been previously documented^23, 24^. Only patients that gave informed consent were included. IRB approvals for this cohort were obtained from NYP/WCMC with reference number 20-05022072 and 1405015116.

#### Non-COVID-19 ICU controls

Twenty-eight patients admitted to NYP hospital ICU between 2014 to 2019 were included as non-COVID-19 ICU controls for the Rep cohort. These patients were suffering from bacterial sepsis ARDS (N = 15), influenza ARDS (N = 4), and influenza pneumonia (N = 9). The patient recruitment, medical history, and sampling procedures for the non-COVID ICU control cohort are the same as those described for the Rep cohort.

### Sengenics assay description and data pre-processing

The Disc cohort samples were processed at Qatar Biomedical Research Institution (QBRI) for KREX immunoproteomics. The Rep cohort samples were processed at the Sengenics facility in Kuala Lumpur, Malaysia. Samples of Disc cohort and controls were analyzed for antigen-specific autoantibodies using Immunome protein arrays (Sengenics), developed using KoRectly Expressed (KREX) technology to provide a high-throughput immunoassay based on correctly folded, full length and functional recombinant human proteins expressed in insect cells, thereby displaying a full repertoire continuous and discontinuous epitopes for autoantibody binding^25, 26^. The Immunome arrays contain more than 1,600 human antigens, enriched for kinases, signaling molecules, cytokines, interleukins, chemokines, as well as known autoimmune- and cancer antigens. Plasma samples of Rep cohort and non-COVID-19 ICU control patients were then processed for autoantibodies on a custom array containing a subset of 1,318 human proteins (Sengenics).

Samples were viral-inactivated in 10% Triton X-100 for 2 hours at room temperature. Samples were then diluted in Serum Albumin Buffer (SAB) at optimized dilution (50-fold dilution). Microarray slides were prepared in four-well plates slide. Samples including controls were randomized and applied to the microarray slides for 2 hours and samples’ IgGs were then detected by secondary Cy3-labeled IgG antibodies. Slides were scanned at a fixed gain setting using the Agilent G4600AD fluorescence microarray scanner generating a 16-bit TIFF file. A visual quality control check was conducted and any array showing spot merging or other artefacts were re-assayed. A GAL (GenePix Array List) file containing information regarding the location and identity of all probed spots was used to aid with image analysis. Automatic extraction and quantification of each spot was performed using GenePix Pro 7 software (Molecular Devices) yielding the median foreground and local background pixel intensities for each spot.

Biotinylated human IgG (detected by fluorescently labelled secondary antibody) and biotinylated human anti-IgG (detected only when plasma or serum is added to the slide) were used as positive controls to assess assay integrity. Extrapolated data was then filtered, normalized and transformed as follows: Briefly, the median background pixel intensity (RFU) was subtracted from the median foreground pixel intensity (RFU) for each antigen to give the median net intensity per spot (RFU); CVs were calculated for each antigen based on the quadruplicate technical replica spots for each antigen on a given array, any antigens with CV above 20% were flagged and outlier spots removed, providing that at least two valid values remained; net intensity values for each antigen in a given sample were calculated as the mean of the net intensity values for technical replica spots on that array; and data was normalised across replica arrays based on the Cy3-BSA controls as previously described^27^. Z-scores were then calculated by subtracting the overall mean antigen intensity (within a single sample) from the net intensity data for each antigen in that sample, and dividing that result by the standard deviation of all of the measured net intensities in that sample, according to the formula: **z = (x – μ) / σ** where **x** is the net intensity of an antigen in a given sample, m is the mean net intensity calculated across all antigens in that sample, and s is the standard deviation of the net intensities for all antigens in that sample. All downstream statistical analysis was done based on the calculated z-scores.

### Sequence identity and antigen specificity analysis for selected proteins

We needed to be cautious in directly comparing the results across different antigens on the arrays because the autoantigen-autoantibody response is not always linear and is an indirect way of prediction of protein concentrations. Since it can depend amongst others on both B cell activation and sequence identity among proteins that express similar antigen epitopes. To check this latter possibility, we selected two proteins (SPANXN4 and STK25) that showed the highest autoantibody alterations to perform their sequence alignment and antigen specificity analysis. Uniprot BLASTP program was used to compare proteins sequences. All human and viral protein sequences with more than 50% sequence similarity were aligned for epitope mapping to determine whether the evaluated RFU values were specific to the protein of interest or could be derived from highly homologous epitopes on other proteins.

### Protein pathways prediction

The assingment of KREX array proteins to functional KEGG categories and their hierarchical organisation was displayed by using Paver, a software for the visualization of Voronoi Treemaps^28^. Any main category is displayed in different colors. The cell sizes were calculated according the signal intensity of the proteins immunofluorescence (highly fluorescent signals give larger cells). Functional Enrichment Analysis was performed to identify biological functions that were over-represented in differentially expressed proteins with a p-value less than 0.05. Differentially expressed proteins, both up-regulated and down-regulated, were used separately as proteins of interest and the proteins detected from all probes were used as the background set. The proteins were further annotated using KEGG- and WIKI-Pathways data prior to performing Fisher’s exact test to determine pathways in which the proteins of interest were significantly over-represented. This analysis was performed on R 3.6.2 using clusterProfiler 3.14.3. GOSemSim was used to eliminate redundant GO-BP results. Only significantly over-represented pathways with a p-value less than 0.05 (-log10 p-value cut-off 1.3) are shown.

### Statistical analysis

Proteins are reported using the symbols of the genes that encode them to offer a clear and uniform nomenclature. Autoantibody response, measured as relative fluorescence units (RFU), was normalized to calculate z-score. Statistical analysis was performed using R (version 4.1.0) and rstudio (version 1.4.1717). Two kinds of inferal statistical tests were performed to test the hypotheses of whether a given autoantibody was differentially expressed in COVID cases versus controls. First, the means between cases and controls were compared using a linear model, using the z-scored autoantibody responses as dependant variables and the COVID state as independent variable (coded as 0=controls and 1=cases). Note that this approach is equivalent to conducting an unrelated T-test and that the effect size of the linear model matches the estimated difference of the means in a T-test. Second, binarized autoantibody responses were tested against cases versus controls using Fisher’s exact test. The cutoff for binarization of the autoimmune response was set to one. As the response is z-scored, this means that all samples with an RFU score above one standard deviation from the mean were considered as being positive for the respective autoantibody whereas all other were considered negative. The Disc and Rep cohorts were analyzed separedly and then merged. Therefore three sets of p-values were obtained for each of the continous and binarized trait analyses.

Following comparisons were made:

- Differential autoantibody response analysis: COVID-19 cases versus controls for
  - Forty-nine Disc COVID-19 patients vs. fourty-eight controls.
  - Forty-eight Rep COVID-19 patients vs. twenty-eight non-COVID-19 ICU controls.
  - Combined ninety-seven COVID-19 vs. seventy-six controls from both Disc and Rep cohorts.
- Pearson’s correlation analysis of fifteen Disc cohort patients sampled at the time of ICU admission (T1) and six weeks follow-up (T2).
- Principal component analysis and pearson’s correlation analysis to compare two cohorts.

## Results

### Study design and cohort specific information

In the present work, two ethnically independent cohorts of COVID-19 patients were evaluated for their autoimmune response (total IgG-response) against 1,318 naturally folded human proteins (antigens). The Disc cohort was recruited at ICU of HMC in Doha, Qatar and included 49 COVID-19 cases and 48 healthy controls, majority of whom were male. A second cohort was recruited from the ICU of NYP Hospital, USA, which included 48 COVID-19 cases and 28 control patients and served as a Rep study. In addition, patients who were admitted to the NYP ICU and had infectious diseases other than COVID-19, such as bacterial sepsis ARDS or H1N1 pneumonia, were included as controls for the Rep cohort. Because of the special composition of the cohorts, we were able to specifically look for COVID-19-related autoantibody signals compared with healthy-baseline- and general infection-baseline-titres. A combined analysis (discovery and replication) allowed stringent COVID-19-specific autoimmune responses to be monitored. Table 1 summarises the demographic and status-specific information of the study cohorts.

**Table 1:** Summary metadata of COVID-19 case and control cohorts

| Study Specification | Condition | Discovery Cohort (Disc) | Replication Cohort (Rep) |
| --- | --- | --- | --- |
| Cohort Size | Control | 48 | 28 |
|  | COVID-19 | 49 | 48 |
|  | COVID-19 Follow-up | 15 |  |
| Gender (Male, Female) | Control | 46 M + 2 F | 19 M + 9 F |
|  | COVID-19 | 48 M + 1 F | 40 M + 8 F |
|  | COVID-19 Follow-up | 15 M |  |
| Age (IQR) | Control | 38 (9) | 66 (26) |
|  | COVID-19 | 47 (20) | 57 (22) |
|  | COVID-19 Follow-up | 49 (11) |  |
| BMI (mean (IQR)) | Control | 25.3 (9.2) | 25.9 (9.4) |
|  | COVID-19 | 29.9 (6.2) | 28.9 (7.3) |
|  | COVID-19 Follow-up | 28.7 (6.4) |  |
| Sampling Time (media days (IQR)) | COVID-19 | 5 (3) days after ICU admission | 6 (6) days after ICU admission |
|  | COVID-19 Follow-up | Six weeks after recovery |  |
| Status Controls |  | Healthy | ICU Bacterial ARDS & Pneumonia, ICU H1N1 ARDS & Pneumonia |
| Status COVID-19 |  | Severe | Severe, ARDS, Pneumonia |
| Hospital |  | ICU, HMC, Qatar | ICU, New York, USA |
| Ethnicity (%) |  | South Asian (69), Middle East and North Africa (MENA) (25). Other (6) | White (31), Asian (10), African (9), Other/Unspecified (50) |
| Matrix, Tubes, and Virus Inactivation |  | Serum, non-EDTA coated, viral inactivation using 10% Triton X100 | Plasma, EDTA coated, viral inactivation using 10% Triton X100 |

### General autoantibody response in healthy and COVID-19 patients

To discover functional IgG-related autoantibodies that could influence COVID-19 predictions and/or outcomes, we used the KREX high-throughput autoantibody assay technology that includes a variety of known human-autoantigens such as cancer-, kinase-, interleukins-, cytokine, ribonuclear transcription and signaling-proteins^29^. Total IgG autoantibody responses were quantified for 1,600 proteins in the Discovery Cohort and for a subset of 1,318 proteins in the Replication Cohort. However, to increase stringency and reduce complexity, only the 1,318 overlapping proteins were subsequently used in the analysis pipeline. The majority of antigens on the array are found in the cytoplasm, nucleus, or cell membrane, but there are also proteins from the mitochondria, endoplasmic reticulum, and cytoskeleton.

The KREX assay reports RFU values for autoantigen-specific autoantibody binding, with linearity over 6 orders of magnitude and with a detection limit in the pg/ml range. These measured RFU values correlate directly with the antigen-specific IgG autoantibody titres, since ligand binding theory shows that the measured signal on-array is linearly proportional to autoantibody concentration. Thus, a higher RFU value for a specific autoantibody-autoantigen interaction indicates a higher autoantibody titre, whilst a higher antibody titer in turn implies a higher autoantigen concentration (or repeated exposure to the autoantigen), accepting that this latter correlation is non-linear. In a first overview, the general intensity distributions were calculated based on the mean autoantibody-antigen titers across all samples and further examined using KEGG-Brite-based Voronoi treemaps using the replication cohort as an example (Figure 1). Approximately 1,150 of the 1,318 proteins could be assigned to the annotation, with the relative size of each cell on the Voronoi treemaps reflecting the observed autoantibody response against that protein (Figure 1 left). Nearly all proteins showed a total IgG AB-signal in the cases and the corresponding controls, the latter represents the natural autoimmunity or the healthy repertoire of autoantibodies. In Figure 1 right, the corresponding pathways are summarized in different colors, with most proteins belonging to the MAPK pathway (light blue), followed by transcription factors (green), chromosomal proteins (green), ribosomes (all blue), and metabolic proteins (yellow). A few proteins belong to the cell cycle (red), chemokines (cyan) or cancer (black). The 10 highest autoantibody titers were found against RBPJ, TPM1, TACC1, KRT19, PTPN20, TBCB, KRT15, AFF4, HSPD1, and CBFA2T3, many of these are structure related proteins. The 10 proteins with the lowest titers were AIF1, IL18, NCK1, COMMD3, NEK11, TGFBR2, SLA, PKM, MAPK6, and MLKL, many of which are cytoplasmic proteins involved in phosphorylation.

**Figure 1:**
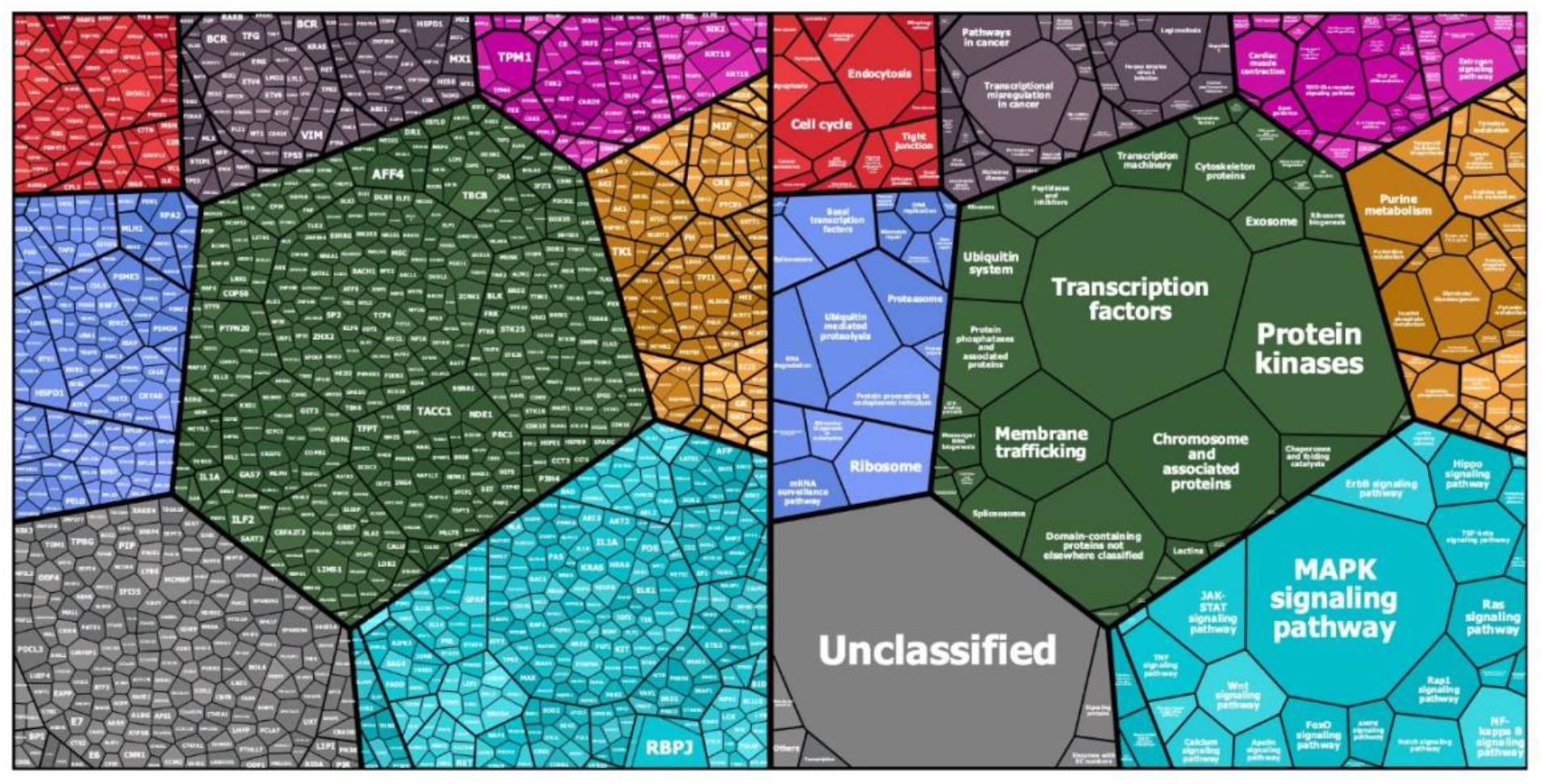
Mapping of KREX Array proteins to KEGG categories (KEGG Pathway and KEGG Brite): Protein symbols and median fluorescent antibody signals (treemap cell size) are represented according their KEGG category assignment (www.kegg.jp; accessed on 14.Nov.2021). The other main categories are defined as cellular processes (top left - red), Human diseases (top middle - greyish purple), Organismal systems (to right - magenta), Genetic information processing (left - blue), Brite protein families (center - dark green), metabolism (right - orange), environmental information processing (bottom right - cyan). Unmapped proteins are considered as “Not included in Pathway or Brite” (bottom left - grey).

### Relative autoantibody response in the Disc cohort revealed significantly higher level of SPANXN4 and ATF4

To examine the effects of SARS-CoV-2 infection on the autoantibody response, we first performed a differential expression analysis in Disc cohort between COVID-19 cases and healthy controls using T-test. Autoantibody responses of fifty-seven proteins were altered significantly (T-test p-value ≤ 0.05) (Supplementary file sheet 3). Autoantibody responses in COVID-19 patients were increased for forty proteins, while decreased for seventeen proteins (Figure 2A). The most elevated autoantibody responses in COVID-19 patients were against ATF4 (effect size (beta) = 3.32 SD; T-test p-value ≤ 0.001) and the sperm protein associated with the nucleus on the X chromosome N4 (SPANXN4) (effect size (beta) = 3.32 SD; T-test p-value ≤ 0.001). The latter is also known as spermiogenesis-related protein and belongs to the family of cancer/testis-associated proteins (CTAs)^30^.

**Figure 2:**
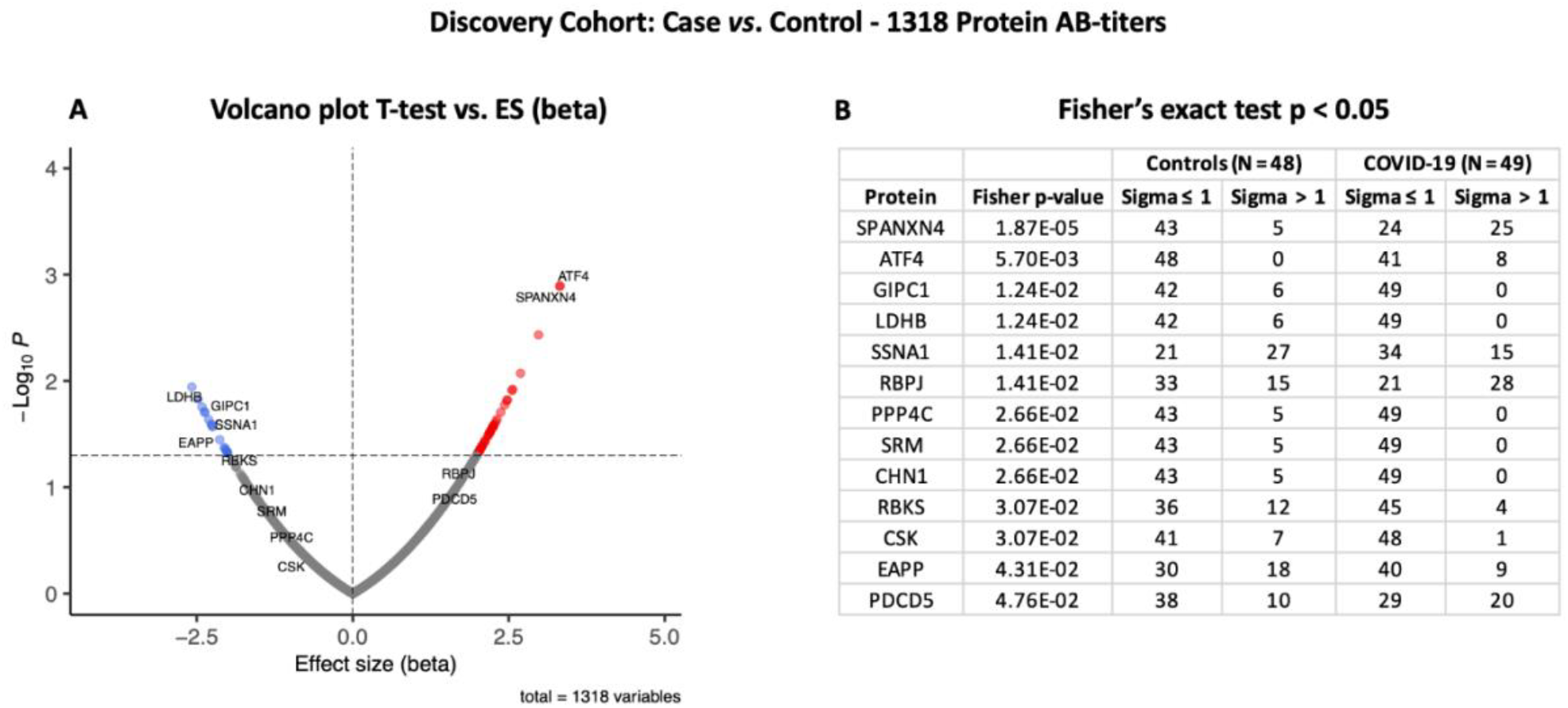
Differential protein autoantibody response analysis of COVID-19 Discovery cohort performed using T-test (A) and Fisher’s exact test (B). A) Volcano graph of 1,318 proteins compares COVID-19 case (n = 49) vs. healthy controls (n = 48). Red dots represent proteins with an elevated autoantibody response, while blue dots represent proteins with a lower autoantibody response in COVID-19 patients. Proteins with Fisher’s test p-value ≤ 0.05 are labelled in the volcano graph. B) Table on Fisher’s exact statistics comparing subjects (numbers) of COVID-19 (n = 49) and the control (n = 48) groups for only thirteen proteins that showed significantly altered (p-value ≤ 0.05) autoantibody responses at sigma > 1.

We then conducted an analysis using binarized autoimmune response, assuming that all samples with an autoimmune response that exceeds the mean by one s.d. as positive and all others as negative (Supplementary file sheet 4). Using Fisher’s exact test, we found twenty-five COVID-19 patients had higher RFU values for SPANXN4 compared to only five in controls (Fisher’s test p-value ≤ 0.0001) (Figure 2B). Autoantibodies against ATF4, recombining signal binding protein J (RBPJ), and programmed cell death 5 (PDCD5) were also significantly elevated (Fisher’s test p-value ≤ 0.05) in the COVID-19 patients. Only the binarized SPANXN4 association reaches the most stringent Bonferroni significance level, that is p < 0.05 / number of proteins = 1,318. In the control group EAPP, SSNA1, and LDHB proteins showed higher autoantibody responses than the cases.

### Autoantibody response in the Disc Follow-up cohort confirmed high levels against SPANXN4 and other proteins in COVID-19 patients

Following the initial blood sample collection at the time of ICU admission, follow-up samples were collected from fifteen patients at six weeks after recovery from COVID-19. For several proteins, a strong correlation (Pearson’s r^2^ ≥ 0.69) was observed between the autoantibody responses at the two sampling time points (Figure 3A). Autoantibody responses against several proteins, including SPANXN4, STK25, TRAF3IP1, AMOTL2, PSMD4, and PPP1R2P9 remained highly elevated (p ≤ 0.05) at 6 weeks post-recovery follow-up. Particularly, autoantibody responses against SPANXN4 (Figure 3B) stayed elevated at both initial (T1) and follow-up (T2) time points. These observations reveal that SPANXN4 autoantibody responses remain elevated for extended periods, suggesting potential association with chronic health issues.

**Figure 3:**
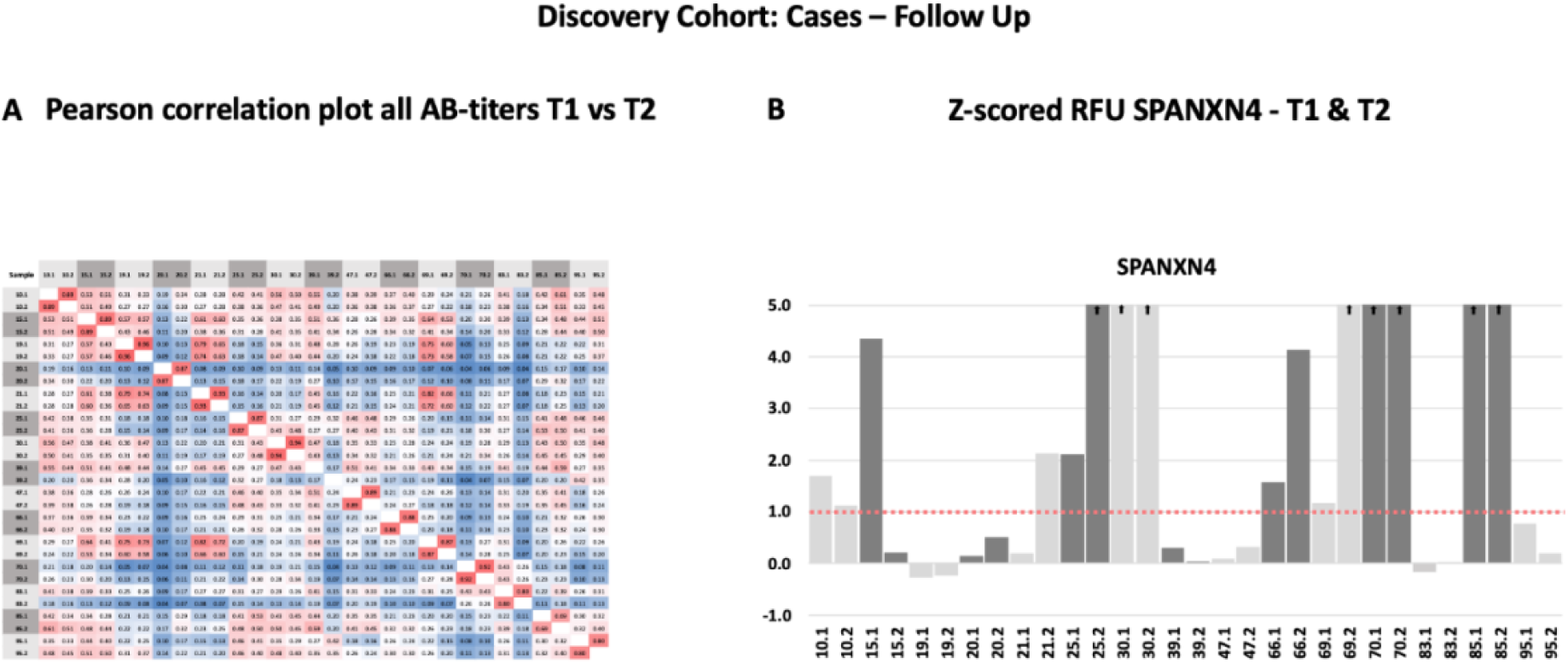
Autoantibody response in the Discovery cohort (T1 = sampling during ICU admission) and follow-up patients (T2 = sampling after recovery). A) Spearman rank correlation analysis of fifteen Discovery cohort samples collected at two different time points show strong correlation (r ≥ 0.69) of autoantibody responses for the proteins. B) The histogram of Discovery cohort samples shows that the z-score RFU of SPANXN4 protein remains elevated in many patients even after COVID-19 recovery.

### Relative autoantibody response in the Rep cohort confirms the trend in the Disc cohort

Autoantibody response for the Rep cohort (n = 48) was compared with the non-COVID-19 ICU control patients (N = 28) (Figure 4A). Autoantibody responses of twenty-six proteins altered significantly (T-test p-value ≤ 0.05) in the Rep cohort. Based on T-test analysis, the most elevated autoantibody response in Rep COVID-19 cohort was found for PRKD2 and BACH1 proteins, which are known for their roles in male reproductive tract development (PRKD2)^31^ and spermatogenesis (BACH1)^32^. Autoantibody response for SPANXN4 was also higher (effect size (beta) = 1.61) in the Rep COVID-19 patients, albeit p-value was slightly higher than 0.05.

**Figure 4:**
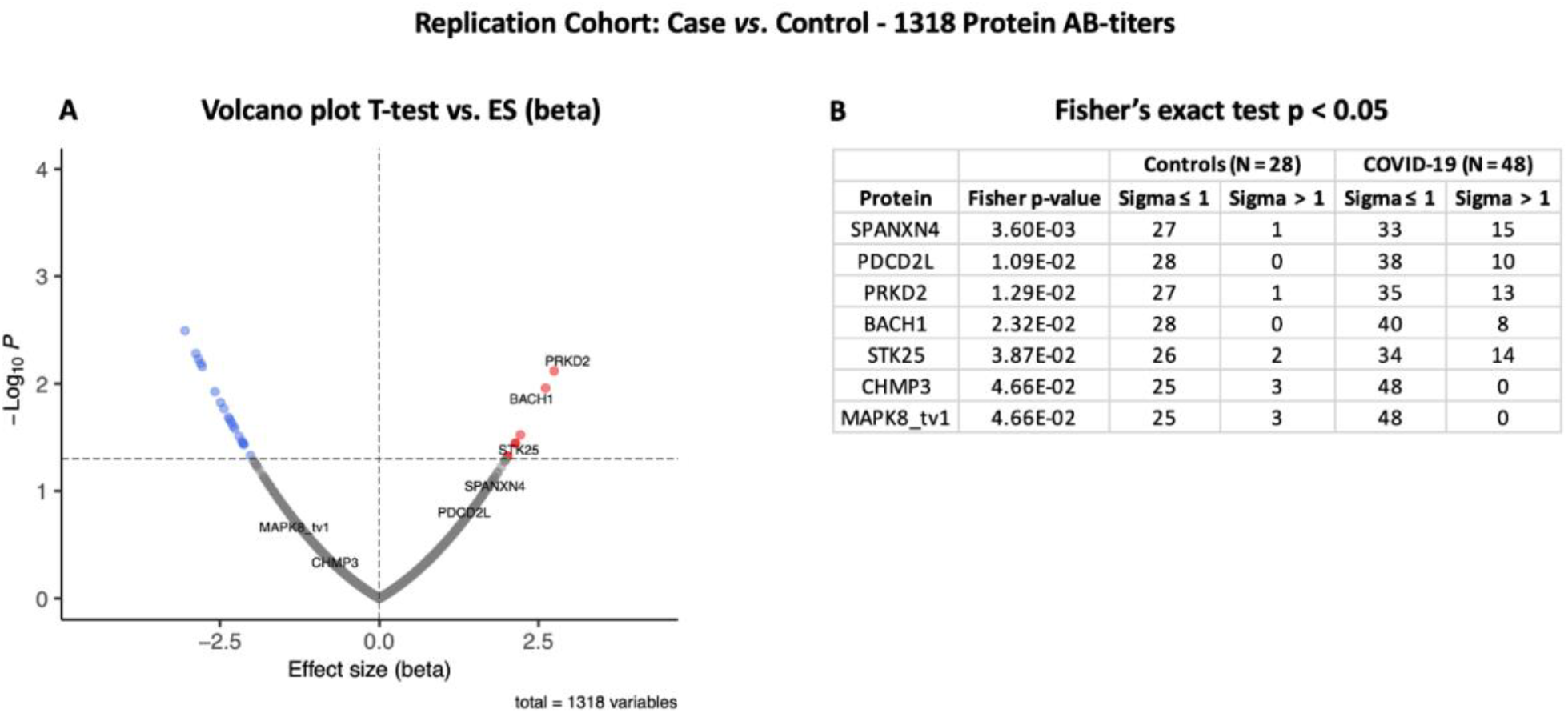
Differential protein autoantibody response analysis of COVID-19 Replication cohort performed using T-test (A) and Fisher’s exact test (B). A) Volcano graph of 1,318 proteins compares COVID-19 cases (n = 48) vs. non-COVID-19 ICU controls (n = 28). Red dots represent proteins with a high autoantibody response, while blue dots represent proteins with a low autoantibody response in COVID-19 positive patients. Only proteins with Fisher’s test p-value ≤ 0.05 are labelled in the volcano graph. B) Table on Fisher’s exact statistics comparing subjects (numbers) of COVID-19 (n = 49) and the control (n = 48) groups for only thirteen proteins that showed significantly altered (p-value ≤ 0.05) autoantibody responses at sigma > 1.

However, Fisher’s exact test indicated that autoantibody response to SPANXN4 remained the highest (Fisher’s test p-value = 0.0036) (Figure 4B). At sigma 1, fifteen COVID-19 patients had higher RFU values for SPANXN4 compared to only one in controls. SPANXN4 can therefore be considered fully replicated under the highest standards of a discovery-replication design. PDCD2L, PRKD2, and STK25 showed also higher autoantibody responses in COVID-19 patients (Fisher’s test p-value ≤ 0.05).

### Analysis of combined cohorts supports that SPANXN4 and STK25 are significantly elevated in COVID-19 patients independent of sampling martix or patients ethicity

Principal components analysis (PCA) of protein RFU data from the two cohorts demonstrated strong overlap between COVID-19 samples and the two cohorts did not separate into discrete clusters (Figure 5). Pearson’s correlation analysis revealed that the autoantibody responses of the two cohorts have high correlation (r^2^ = 0.73).

**Figure 5:**
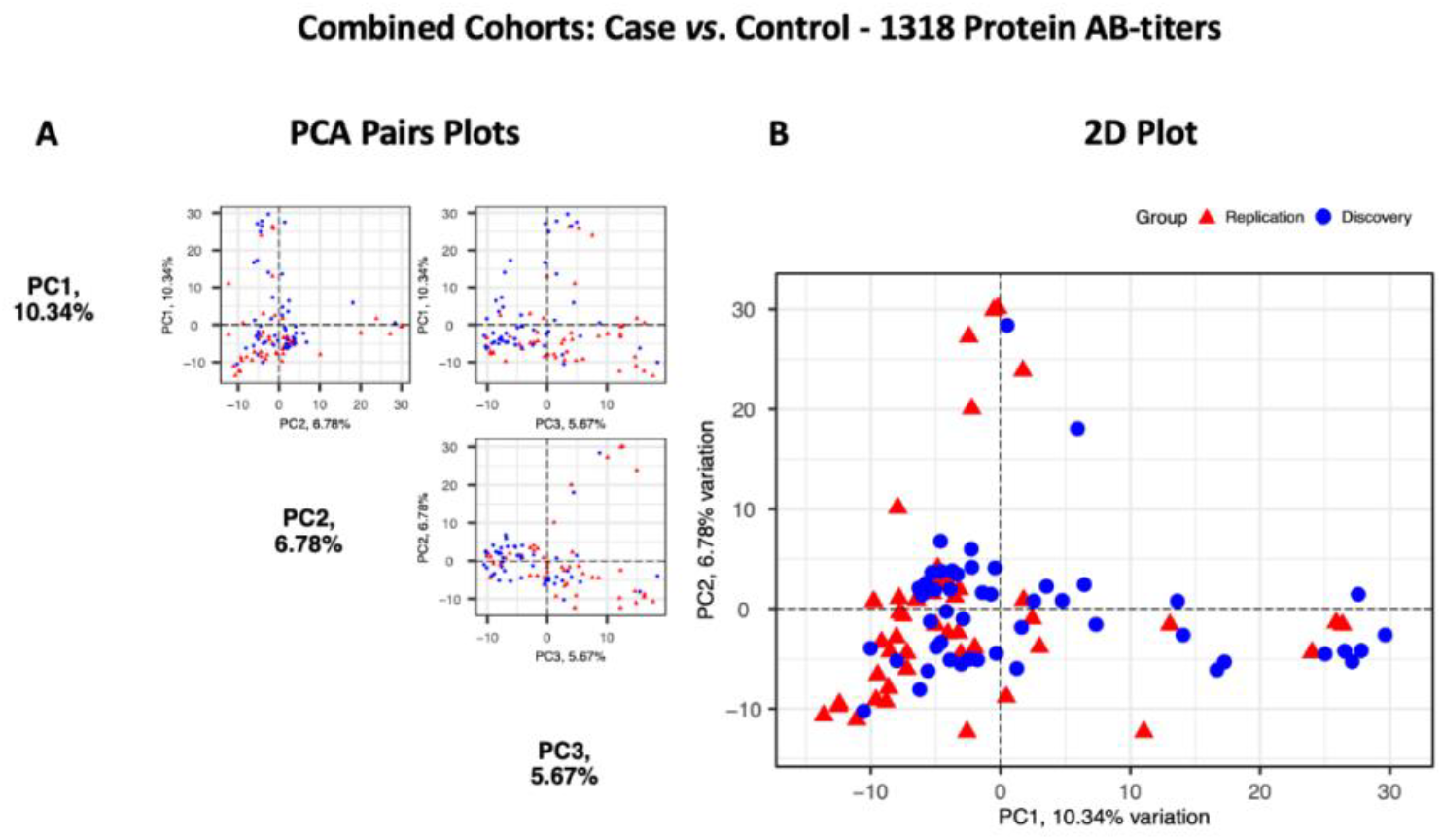
Principal Components Analysis of the Discovery (n = 49, blue circles) and the Replication cohorts (n = 48, red triangles). Each point represents a sample. A) PCA pair plot compares PC1 to PC3. The proportion of variance explained in our cohorts by each PC is shown in parentheses on the axis labels. B) PCA 2D plot with PC1 and PC2, which together describe 17.12% diversity between the cohorts.

At third stage, we combine data from both the Disc and Rep cohorts (n = 97) and compared them with combined controls (n = 76). Case vs. control analysis revealed that autoantibody responses against fifty-six proteins were significantly altered: 35 autoantibodies with increased and 21 autoantibodies with decreased responses (T-test p ≤ 0.05) (Figure 6A). SPANXN4, ATF4, STK25, and PRKD2 were the proteins with the highest effect size (beta). In total forty patients had SPANXN4 RFU higher than 1 sigma value (Fisher’s exact test p-value ≤ 0.0001) in the combined COVID-19 cohorts compared with the six patients only in controls (Figure 6B).

**Figure 6:**
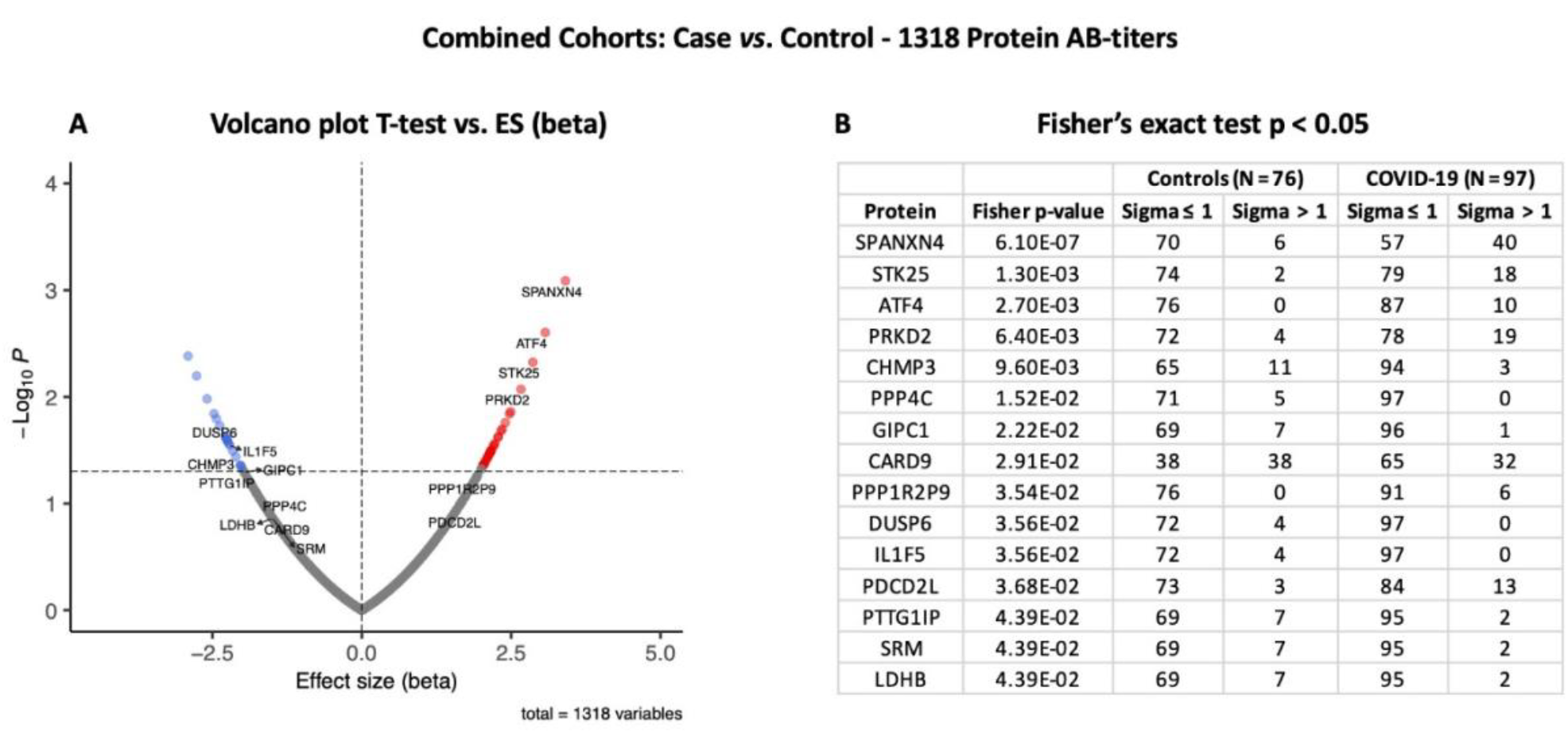
Differential protein autoantibody response analysis of Combined Cohorts performed using T-test (A) and Fisher’s exact test (B). A) Volcano graph of 1,318 AB-protein titers compares COVID-19 cases (n = 97) vs. controls (n = 76). Red dots represent proteins with a high autoantibody response, while blue dots represent proteins with a low autoantibody response in COVID-19 positive patients. Only proteins with Fisher’s test p-value ≤ 0.05 are labelled in the volcano graph. B) Table on Fisher’s exact statistics comparing subjects (numbers) of COVID-19 (n = 49) and the control (n = 48) groups for only thirteen proteins that showed significantly altered (p-value ≤ 0.05) autoantibody responses at sigma > 1.

Furthermore, the autoantibody responses, expressed as RFU z-score for fifty-six proteins that differed significantly between the study groups are shown in Figure 7A. The heatmap shows that most of the proteins display similar pattern of autoantibody ratios across the study cohorts. These analyses demonstrate that our autoantibody response data are highly reproducible despite differences in population ethnicity, different laboratories, and sampling materials (serum vs. plasma in Disc vs. Rep cohorts, respectively).

**Figure 7:**
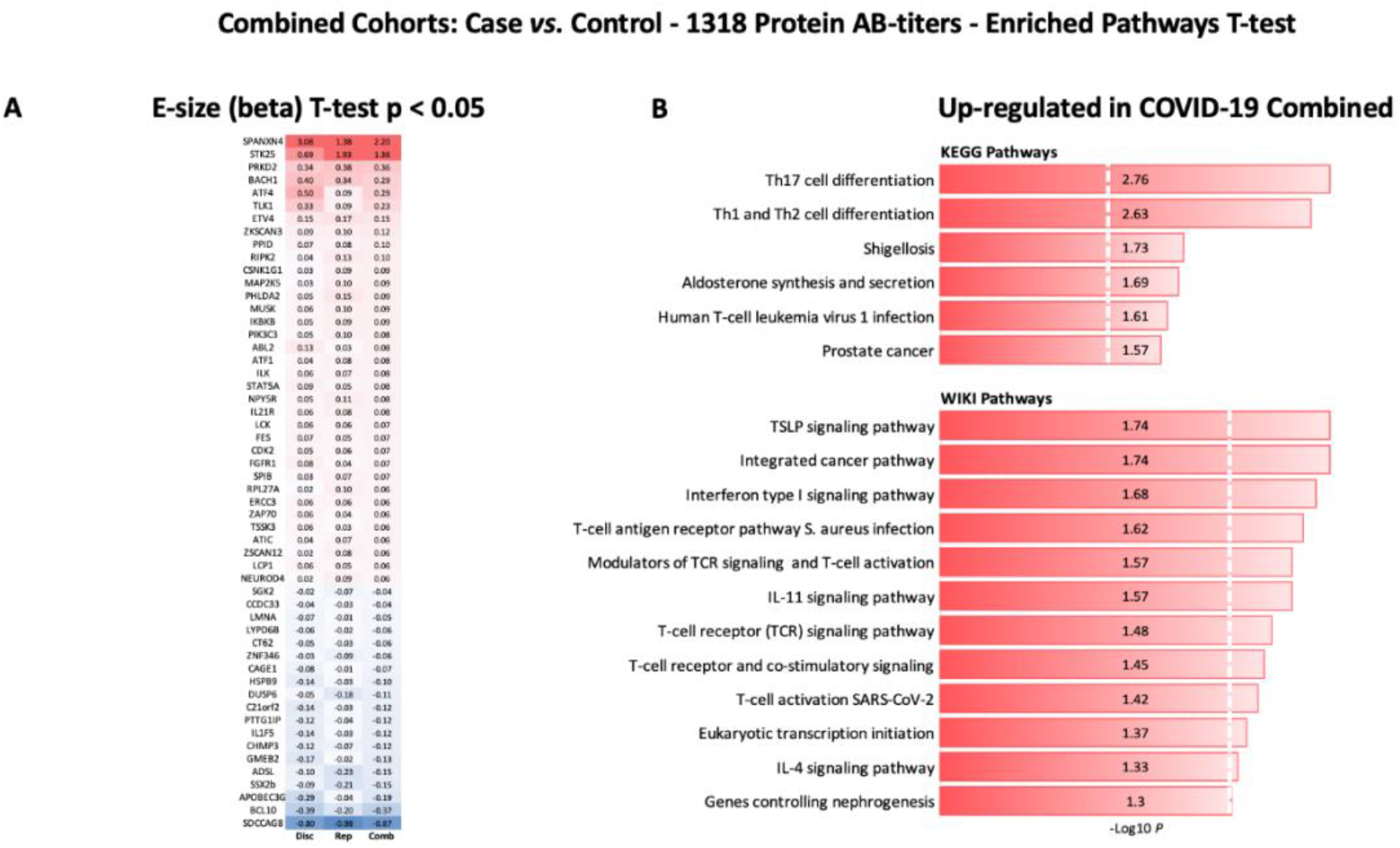
Heatmap of autoantibody response pattern among different cohorts (A) and bar plots for pathways analysis. A) Heatmap of relative estimates (case vs. control) of autoantibody responses to fifty-six proteins in the Discovery, Replication, and Combined cohorts. Only proteins with significantly altered autoantibody responses were selected. Red color indicates higher and blue color indicates lower autoantibody responses against the proteins. B) KEGG and WIKI pathways analysis presented as bar-plot shows overactivated pathways in COVID-19 patients. Only pathways with T-test p-value ≤ 0.05 are presented in the bar-plot.

### Protein pathways analysis uncovered up-regulated immune pathways in COVID-19 patients

KEGG and WIKI pathways analysis was performed to identify the functional contribution of autoantibodies targeted proteins in cellular processes and immune-inflammatory systems. Pathways associated with T helper cells (Th1, Th2, and Th17) differentiation, bacterial/viral infections, stress hormones release, and prostate cancer were upregulated in COVID-19 patients (Figure 7B). WIKI pathways were also activated for host immunity and interferon signaling, including T cell activation for SARS-CoV-2 and *Staphylococcus aureus* infections (Figure 7B).

### SPANXN4 and STK25 share sequence identity with SPANX- and STK-family proteins but showed unique AB-titers in COVID-19 patients

In order to check cross-reactivities, sequence homology and antigen specificity analysis were performed for SPANXN4 and STK25 against human and viral protein databases. Only few proteins appeared to have more than 50% sequence identity with our target proteins ((SPANXN4 with SPANXN1, 2, 3, and 5) and (STK25 with STK 3, 4, 24, and 26)) (Figure 8 and 9). However, many of these homologous proteins were also part of our KREX immunome panel but did not show any significant changes, which means that the observed RFUs are highly specific against the targeted proteins.

**Figure 8:**
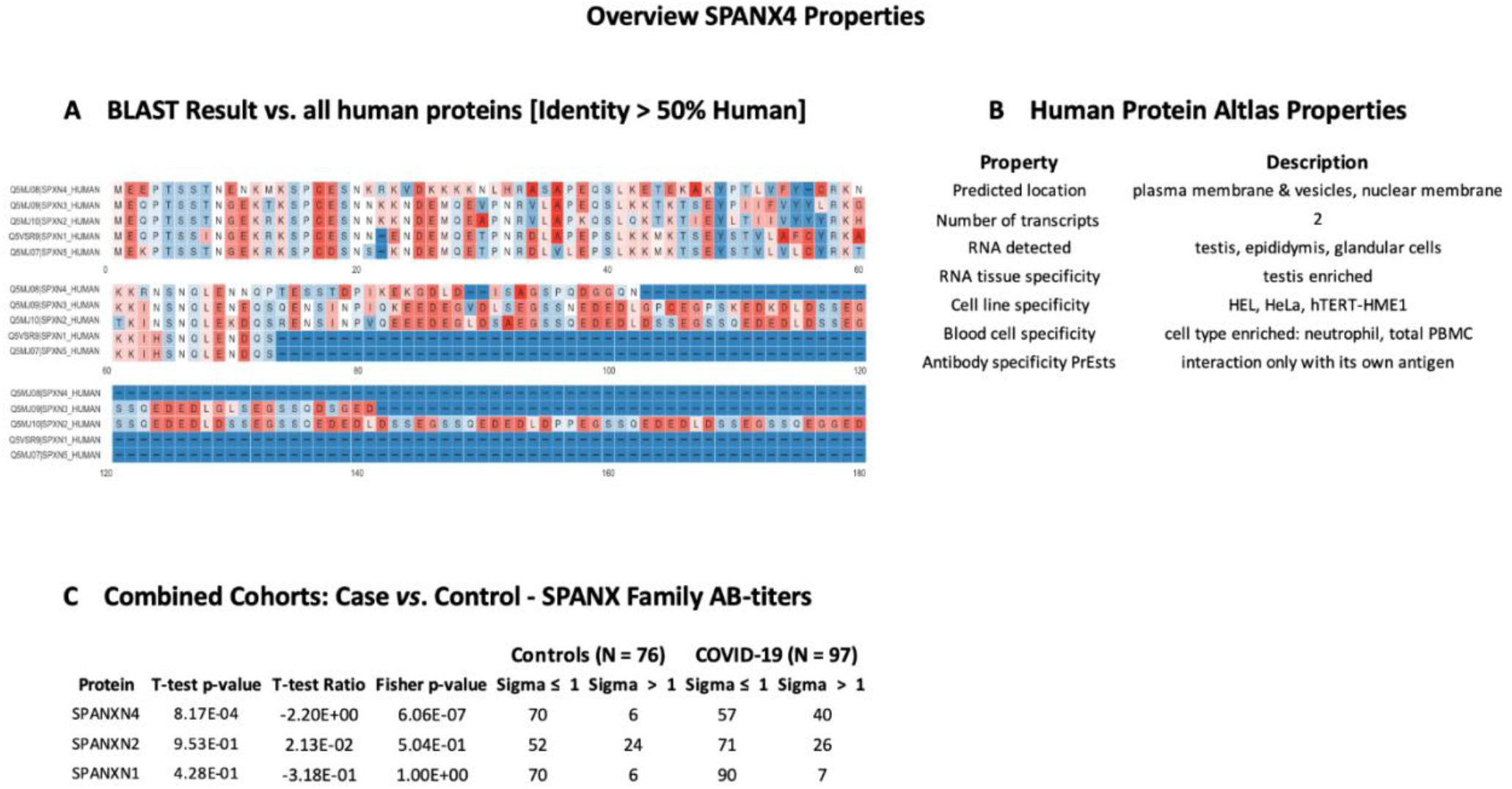
SPANXN4 protein sequence identity and antigen specificity analysis. A) Sequence alignment of proteins showing ≥ 50% identity with SPANXN4. B) Summary of properties of SPANXN4 protein from Human protein atlas. C) Autoantibody responses to SPANXN family proteins with high sequence identity to SPANXN4 that were part of the KREX immunome panel.

**Figure 9:**
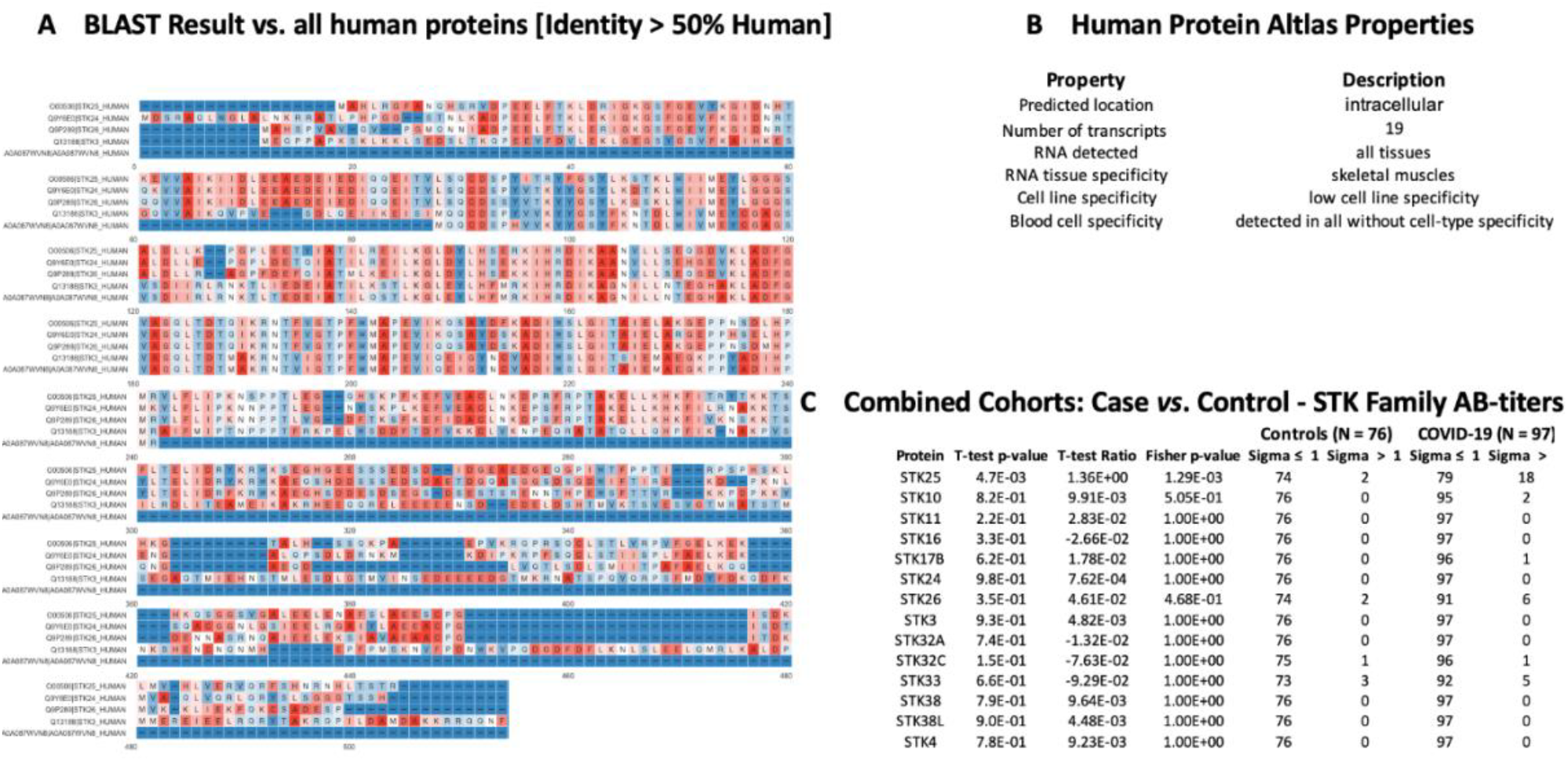
STK25 protein sequence identity and antigen specificity analysis. A) Sequence alignment of proteins showing ≥ 50% identity with STK25. B) Summary of properties of STK25 protein from Human Protein Atlas. C) Autoantibody responses to STK family proteins with high sequence identity to SKT25 that were part of the KREX immunome panel.

## Discussion

In the current COVID-19 pandemic, there is increasing interest globally in understanding the underlying immunology of COVID-19, as well as revealing new health issues arising from COVID-19 complications. Several papers have described the existence and cross-reactivity of SARS-CoV-2 specific T-cell responses^33, 34, 35, 36^, as well as correlations with male reproductive system and infertility^4, 37^. The present study identified and validated several autoantibody responses by screening two independent cohorts of COVID-19 patients with the KREX immunome protein array. The proteins identified with higher autoantibody responses serve important physiological functions and are strongly associated with various immunological and pathological parameters associated with COVID-19 disease.

The KREX immunome array contains proteins involved in physiological processes such as MAPK signaling, metabolism, transcription, cell cycle, immunity, and cancer-related pathways. Few of the proteins with the highest mean autoantibody response in COVID-19 patients were RBPJ, TPM1, TACC1, KRT19, and PTPN20. These proteins perform a variety of physiological functions in the human body, with many of them being structural proteins involved in tissue damage and repair mechanisms^38^. The presence of a high autoantibody response to these proteins suggests that they are overproduced during a pathological condition, such as cancer or a cardiovascular complication^39, 40^. For example, notch signally protein RBPJ has been associated with COVID-19 pathophysiology and cardiovascular complication^41^. Similarly, keratin family proteins (KRT19 and KRT15) that are responsible for epithelial cell structural integrity are linked to COVID-19 pathogenesis and disease severity^42^. Furthermore, many of these proteins are also involved in male reproductive system physiology and fertility, yet there has been no previous report in COVID-19 patients.

Our Disc cohort reported higher autoantibodies against SPANXN4, ATF4, RBPJ, and PDCD5 proteins compared to the controls. Comparison between COVID-19 baseline (T1) vs. follow-up (T2) samples indicated that SPANXN4 autoantibodies remained elevated at post-recovery stage. Prolonged autoantibody responses may highlight COVID-19 post-acute sequelae by stimulating the humoral immune response in a way that leads to long-term autoantibody production^43^. The diverse variety of proteins linked to a prolonged autoantibody response suggest that SARS-CoV-2 may stimulate autoantibody formation by molecular mimicry^44^, targeting cardiolipin, cardiolipin-binding proteins, platelet factor 4, prothrombin, and coagulation factors, suggesting their role in coagulopathies, chronic comorbidities and post-infection recovery^45, 46, 47^. We hypothesize that elevated autoimmune antibodies against SPANXN4, STK25, TRAF3IP1, AMOTL2, PSMD4, and PPP1R2P9 might suggest a similar role. However, Dotan et al.^48^ investigated *in-silico* sequence homology of all human proteins with the virus but could not find evidence that any of the proteins mentioned here are part of such a mimicry process. Vice versa, we cannot exclude that the titers might be elevated before the exposure to SARS-CoV-2, due to pre-existing diseases such as cancer or prolonged inflammation.

In contrast, the Rep cohort had higher levels of autoantibody responses to SPANXN4, PDCD2L, PRKD2, and STK25 proteins than the controls. Except for SPANXN4, all other proteins with the high autoantibody response were not significantly elevated between the two cohorts but often showed similar trends. These differences could be attributed to the fact that the control group in the Disc cohort was comprised of healthy volunteers, whereas the control group in the Rep study was comprised of ICU patients suffering from bacterial or viral ARDS, or pneumonia.

When all COVID-19 patients (N = 97) from both cohorts were merged and compared to all controls (N = 76) from both cohorts, the most significant autoantibody responses were observed against SPANXN4, ATF4, STK25, and PRKD2. ATF4 regulates metabolic and redox processes in the human body, and an increased ATF4 response has been observed in previous coronavirus disease^49, 50^. Fischer et al.^18^ suggested that ATF4 also plays role in differentiation of the vas deferens lamina propria layer that helps improve spermatozoa fertilization rate. STK25 and PRKD2 are two important kinases with several physiological roles in our body. However, their role in male reproductive tract physiology is least discussed. A few studies highlight STK25 as androgenic kinase^16, 17^ and PRKD2 role in male reproductive tract development^19^.

SPANXN4 belongs to a protein family called “sperm protein associated with nucleus in the X chromosome” (SPANX) that are essential for motility and fertilization capacity of male-ejaculated spermatozoa^15^. SPANXNs are also known as cancer testis antigens (CTAs) because of their overexpression in tumor tissues, in addition to their normal physiological role in the testis and spermatozoa of healthy males^51^. SPANX proteins are expressed in various regions of sperms, and research has shown that the protein family has relevance to male fertility. Particularly, the presence of ACE2 receptors in testicular tissues suggests that SARS-CoV-2 influences male fertility, but the pathogenesis is not clear. No previous COVID-19 study has mentioned SPANXN4 involvement in male infertility, neither the pathogenesis is explained. Therefore, the autoantibody response measured in the current study may suggest a novel diagnostic and treatment marker for male fertility. Previously, one hepatitis C virus study has shown that SPANXN4 interacts with the virus, potentially increases virus infectivity, albeit no reproductive performance was discussed^52^. Increased levels of autoantibodies against testis-related proteins suggest their role in affecting male reproductive system, and thus declining male fertility in COVID-19. Although several investigations have found that COVID-19 patients have altered seminal parameters and decreased reproductive hormone levels^53^, histological or functional abnormalities in male genital system^54^, damaged blood-testis barrier^55^, and impaired spermatogenesis^56^, the cause of this comorbidity has not yet been investigated, and remains unknown.

Despite their ethnic diversity, which included Middle Eastern, Africans, Caucasians, Asians, and South Asians populations, correlation analysis, hierarchical, and PCA clustering demonstrated that both cohorts shared similarities in autoantibody responses. Therefore, strong correlation (r^2^ = 0.73) between these cohorts demonstrate that autoantibody response data of COVID-19 patients are highly reproducible among different ethnic populations. These findings are consistent with our prior COVID-19 proteomics study that looked at immune-inflammatory markers in five different demographic cohorts (manuscript accepted).

Several previous COVID-19 studies have reported elevated immune-inflammatory responses, including cytokine-storm in COVID-19 patients. Perhaps, we observed relatively elevated albeit non-significant autoantibody responses to immune cytokines like IL1A and IL1B proteins. KEGG and WIKI pathways analysis showed that autoantibody responses to immune proteins activated T cell responses against infection and T helper cell differentiation. Li et al.,^57^ observed that Th17 differentiation and cytokine response pathways play a key role in pathogenesis of COVID-19 and autoimmune diseases. Pathways analysis suggests that many immune cell responses specific to SARS-CoV-2 or bacterial infections may precede chronic inflammatory disorders and the respiratory failure^58^. Furthermore, an abnormal T helper cell response, combined with overactive interferon signaling, promote the differentiation of B cells, which produce autoantibodies and cause autoimmune diseases^59^.

In conclusion, these findings reveal unique autoantibody response against several proteins that play diverse though important function in COVID-19 complications. These observations also highlight the importance of the humoral immune response, as well as numerous other previously unknown immunological pathways in COVID-19 pathogenesis. Particularly, elevated levels of autoantibodies against the testicular tissue specific protein SPANXN4 in both cohorts offer significant evidence of anticipating the protein’s role in COVID-19 associated male reproductive complications. Overall, these finding not only revalidate autoantibody responses against SPANXN4 in COVID-19 but also predict novel pathological associations that may contribute to COVID-19 post-recovery comorbidities. SPANXN family proteins are known as CTAs^60^ that play essential role in spermatogenesis, however, their role in male fertility in the COVID-19 patients is previously unknown.

## Author Contributions

F.S., H.B.A. and O.M.E. designed, conceived, and led the study. M.A.Y.A., V.M.A., and A.M. led the Disc Cohort sample collection, processing, and ethical approvals. H.B.A. and N.V. optimized the assays on Disc Cohort. I.B. and H.B.A. run the assays on Disc samples. K.S. and M.U.S. performed statistics. E.J.S., D.P., H.S., and A.M.K.C. organized and collected samples for Rep Cohort. T-M.T., P.E.M., and J.M.B. run the assays on Rep samples J.B. and F.M. performed annotation and treemap analyses. K.S., F.S., F.M., M.U.S., H.B.A., O.M.E., J.D., J.B., and A.A. interpreted the data. M.U.S., K.S., H.B.A., and F.S. wrote the manuscript. All authors reviewed the manuscript and have read and agreed to the published version of the manuscript.

## Institutional Review Board Statement

The study on Qatar Cohort was conducted according to the Ministry of Public Health (MOPH) guidelines and approved by the Institutional Review Board Research Ethics Committee of the Hamad Medical Corporation (reference MRC-05-003).

## Acknowledgments

We would like to thank all the patients, volunteers, and healthcare co-workers from HMC and NYP/WCMC hospitals, and Rudolph Engelke for providing the R-code for functional enrichment. We also acknowledge the help provided by the Anti-Doping Lab-Qatar (ADLQ) and Qatar Red Crescent (QRC) for recruiting control individuals. The Qatar cohort research was supported by the Proteomics core facility and ID-IDRP project at QBRI. This work was also strongly supported by the Biomedical Research Program at Weill Cornell Medicine in Qatar, a program funded by the Qatar Foundation.

